# The pharmacogenomic assessment of molecular epithelial-mesenchymal transition signatures reveals drug susceptibilities in cancer cell lines

**DOI:** 10.1101/2024.01.16.575190

**Authors:** Alexander J. Ohnmacht, Göksu Avar, Marisa K. Schübel, Thomas J. O’Neill, Daniel Krappmann, Michael P. Menden

## Abstract

The epithelial-mesenchymal transition (EMT) is characterised by the loss of cell-cell adhesion and cell polarity, which is often exploited by cancer cells to adopt a motile, invasive and metastatic phenotype. Whilst EMT is often linked with cancer progression and therapy resistance, strategies for its selective targeting remain limited. In order to address this, we infer EMT states of cancer cell lines from their molecular signatures and use predictive and causal modelling to estimate the effect of EMT on drug susceptibility in high-throughput drug screens. For example, we show that EMT signatures in melanoma cells can predict favourable responses to the HSP90 inhibitor luminespib and demonstrate that epithelial-like melanoma cells can be sensitised to luminespib upon stimulation of EMT by TGF-β. Thus, we provide an analysis that systematically yields a set of potent drugs by exploiting vulnerabilities of cancer cells undergoing EMT, which may pave the way for therapies to target these cells.

## Introduction

The epithelial-mesenchymal transition (EMT) is a cellular process that allows cells to transition between different phenotypic states ^1^. Rather than a switch between two distinct phenotypes, the EMT program describes a dynamic spectrum of phenotypes between epithelial and mesenchymal cells ranging from apical-basal polarity and strong cell-cell contacts to motile and spindle-like characteristics ^2^. EMT is an essential mechanism for embryonic development, wound healing and tumour plasticity, and has been regarded as a hallmark of cancer ^1,3–5^. The invasion of the extracellular matrix by cancer cells undergoing EMT prior to metastasis is accompanied by the loss of the adherens junction protein E-cadherin and upregulation of N-cadherin, vimentin and fibronectin^1,6^. Scores derived from gene expression signatures of these molecular markers are typically used for assessing EMT and its associated cellular processes in cancers ^2,7–10^. Some of these processes can be used to externally stimulate cells to undergo EMT. For example, TGF-β signalling is an established mechanism for inducing EMT ^11^ and thus TGF-β treatment is widely used for external EMT induction *in vitro* ^*12–17*^.

Sparse findings in cancer cell lines and human tumours have reported EMT as a putative drug response biomarker ^7–10^. For example, acquired resistance through EMT has been reported for commonly employed chemotherapeutic agents, e.g. cisplatin and doxorubicin ^18^, and targeted therapies, e.g. EGFR or PI3K inhibitors ^19^. Furthermore, EMT was found to cause intrinsic resistance to KRAS inhibitors in lung cancer ^20^. Although the genetic background has been shown to play an important role in enabling EMT in cancer progression ^10^, it is still unclear to what extent initial cancer drug responses can be attributed to EMT. Thus, we hypothesised that predictive and causal modelling of EMT scores in drug high-throughput screens can assess the role of EMT in cancer drug sensitivity, which may lead to strategies that systematically exploit EMT as a cancer vulnerability.

Here, we first estimated continuous EMT scores based on gene expression profiles of 790 cancer cell lines from 31 cancer types using four different methods ^7,8,10,21^. Consecutively, we benchmarked the contribution of EMT in drug response prediction models and quantitatively estimated the EMT effect with causal inference. For example, we revealed that EMT and its related processes in melanoma robustly predict sensitivity to HSP90 inhibition with luminespib and other HSP90 inhibitors. Indeed, we experimentally demonstrated that stimulating EMT with TGF-β pretreatment can sensitise epithelial melanoma cell lines to luminespib.

## Results

We leveraged a high-throughput drug screen (HTS; **Fig. 1a**) of 790 cancer cell lines across 31 cancer types, which were treated with 544 unique compounds to obtain dose-response curves (**Fig. 1b**) ^22–24^. This was complemented with molecular profiling of cancer cell lines, i.e. somatic mutations, copy number alterations and gene expression (**Fig. 1c**) ^22–24^. For estimating EMT, we derived EMT scores from four established methods that leverage molecular signatures to infer EMT on a continuous spectrum using gene expression data (**Fig. 1c**); these were: Mak *et al*. ^8^, gene set variation analysis ^21^, Tan *et al*. ^7^ and Tagliazucchi and Wiecek *et al*. ^10^ (**Methods**), abbreviated as MAK, GSVA, TAN and TW, respectively (**Supplementary Data 1**). Then, we systematically benchmarked the EMT scores for predicting drug responses across all compounds and cancer types using (1) ablation of the EMT score and (2) causal inference of the EMT effect (**Fig. 1c**; **Methods**). Thereby, the cancer somatic alterations served as background predictors for assessing the EMT-specific component.

**Figure 1:**
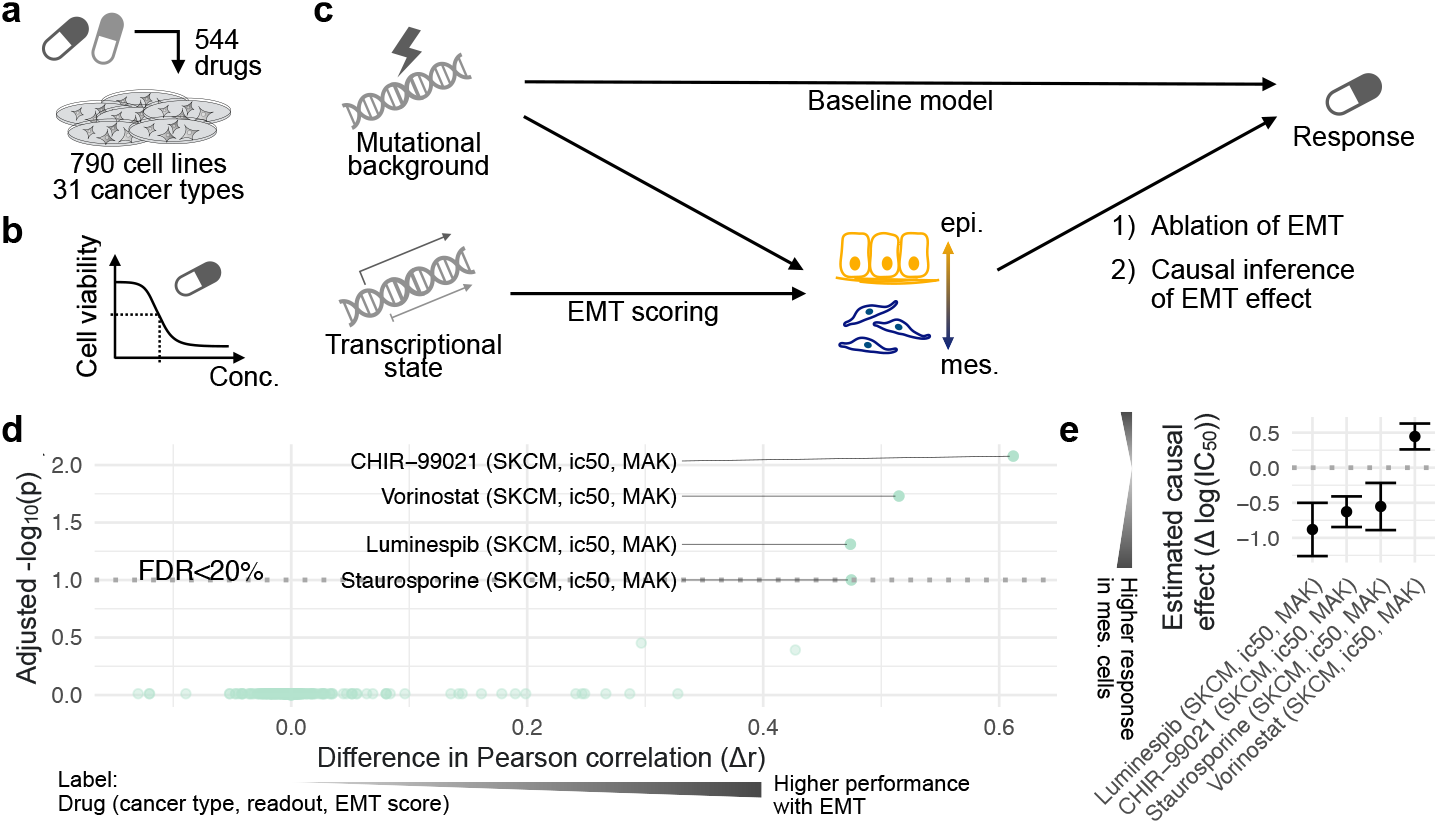
Modelling drug susceptibility in cancer cell lines in the context of EMT. (**a**) The drug high-throughput screen contained 790 cancer cell lines treated with 544 compounds and (**b**) their dose-response curves. (**c**) Molecular profiling of cancer cell lines quantifies their mutational background and the transcriptional state. Shown in the schematic workflow for predictive and causal modelling, the mutational background are baseline features and EMT scores are derived from gene expression data. First, a regression-based ablation study assessed the predictive performances of the drug response prediction model upon excluding EMT from the model. Secondly, the EMT-specific effect on drug susceptibility was estimated by causal inference methods (**Methods**). (**d**) The systematic ablation study in SKCM yielded a set of compounds for which EMT improved the response predictions, showing the adjusted *p*-values of a *t*-test for performance metrics and the difference in Pearson’s correlation Δr. (**e**) The inferred EMT effects on responses to the identified set of compounds with the 95% confidence interval in SKCM is shown.

Exemplifying our method, we leveraged the MAK EMT scores and drug responses quantified by IC_50_ values in skin cutaneous melanoma (SKCM) and identified four inhibitory compounds, for which the full model including EMT significantly outperformed the baseline model (**Fig. 1d**; **Methods**). For example, response to luminespib in SKCM was predicted well by the full model, i.e. leveraging EMT scores and the mutational background with Pearson’s r = 0.50 between actual and predicted IC_50_ values. However, the performance of actual versus predicted IC_50_ dropped to Pearson’s r = 0.02 upon exclusion of the EMT score (Δr = 0.47, *p* = 5.0 ×10^−4^, *t*-test for resampled performance metrics; **Supplementary Fig. 1a**). For the identified compounds, we applied double machine learning in conjunction with causal random forests to estimate the EMT-specific effect on drug responses with a 95% confidence interval (**Fig. 1e**; **Methods**) ^25–28^. Compounds with significantly increased performance and high inferred effect size for multiple EMT scores and both IC_50_ and area under the drug response curve (AUC) suggested a causal component of EMT for determining drug responses (**Supplementary Fig. 1b-f**). This hypothesis was systematically dissected across the remaining cancer types, EMT scores and drug response readouts in the next section.

### Systematic analysis of EMT and its regulators as biomarkers of cancer drug sensitivity

The distributions of MAK, GSVA, TAN and TW scores were predominantly cancer type specific (**Fig. 2a-d**). For example, SKCM cell lines showed a more mesenchymal MAK EMT score, whilst breast cancer (BRCA) and colorectal cancer (COREAD) cell lines displayed rather epithelial MAK EMT scores (**Fig. 2a**), which highlighted the high tissue-specificity of EMT molecular signatures. MAK, TAN and TW scores showed high overall correlations (Pearson’s r > 0.87; **Supplementary Fig. 2a**), which were consistently high within cancer types. GSVA showed lower overall correlations with these scores (Pearson’s r < 0.39; **Supplementary Fig. 2a**) due to normalised scores (**Supplementary Fig. 2b**), but displayed consistently high correlations within cancer types as well (**Supplementary Fig. 2a**).

**Figure 2:**
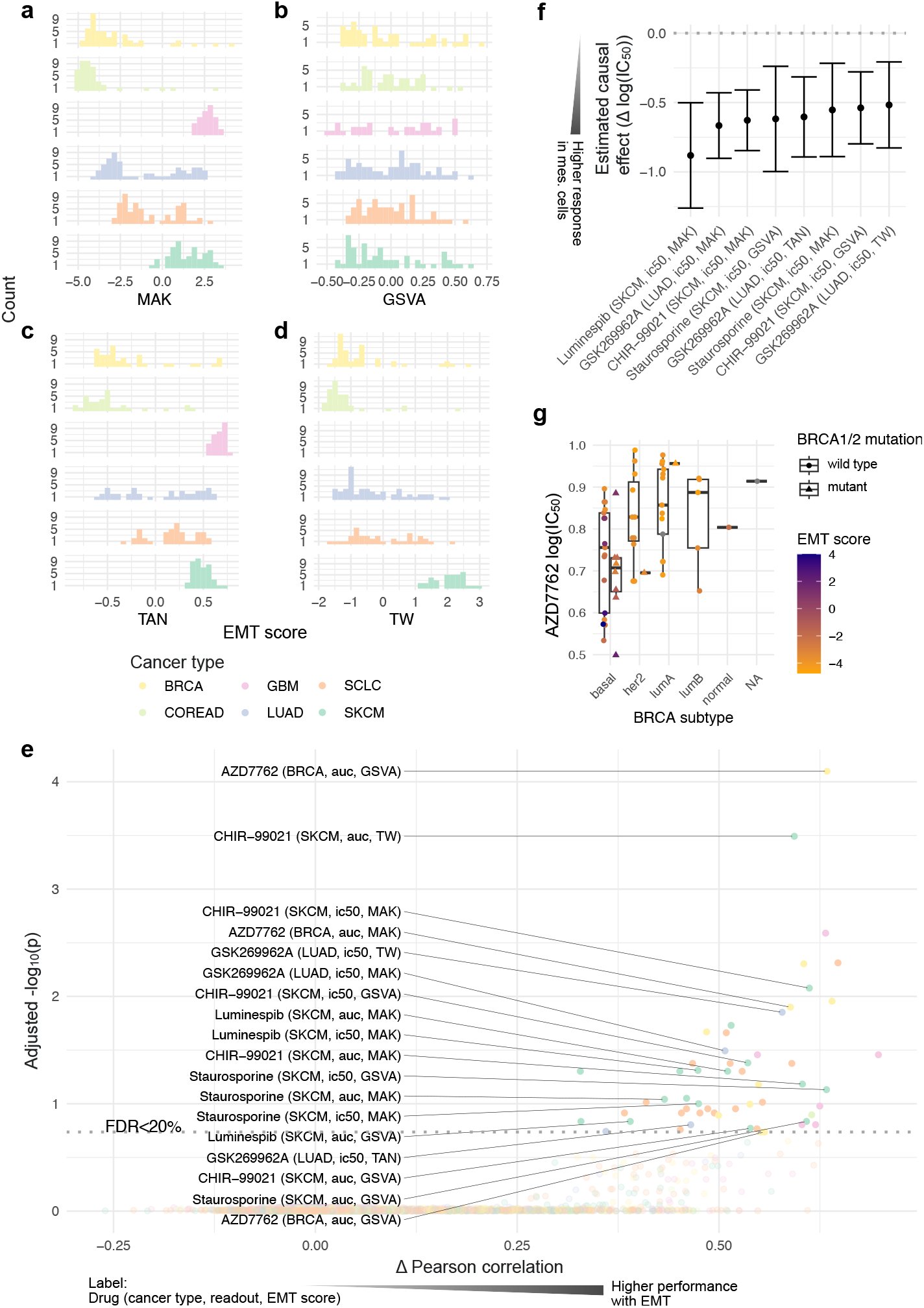
EMT as a predictive component of drug sensitivity. Distributions of (**a**) MAK, (**b**) GSVA, (**c**) TAN and (**d**) TW EMT scores depending on the cancer type are displayed. (**e**) The systematic ablation study demonstrates EMT as a predictor of drug sensitivity in cancer cell lines for four different EMT scores and two response readouts, i.e. IC_50_ and AUC, with six cancer types that showed at least one significant compound (FDR < 0.2). The compounds and cancer types that showed at least 3 significant performance changes are labelled. The horizontal axis represents the difference in mean Pearson’s correlation Δr between predicted and actual IC_50_ or AUC values of the models, including and excluding EMT, whereas the vertical axis measures the significant improvement in the performance over the baseline model using a *t*-test for resampled performance metrics. (**f**) For the IC_50_ prediction models, the estimated EMT effects plus 95% confidence intervals are displayed. (**g**) The boxplot shows the response to CHK1/2 inhibitor AZD7762 depending on BRCA PAM50 subtypes, the MAK EMT score and mutations in *BRCA1/2*. The centre on the boxplot represents the median, while the box illustrates the interquartile range (IQR). The whiskers show a range that is 1.5 times the size of the IQR.

We conducted the benchmark with the outlined modelling strategies (**Fig. 1c**; **Methods**), and recorded its results across all included cancer types, EMT scores, compounds and IC_50_ or AUC (**Supplementary Data 2**; **Methods**). Six cancer types showed at least one significant compound with FDR < 0.2 (**Fig. 2e**; **Methods**). We estimated the EMT effects and confidence intervals for all compounds (**Supplementary Fig. 3a,b**; **Methods**), and further focused on five compounds in three cancer types that consistently showed significant performance gains in at least three models (labelled in **Fig. 2e,f**; **Table 1**; **Methods**). For all of these compounds, mesenchymal cells showed higher drug responses than epithelial cells (**Fig. 2f**).

**Table 1:**
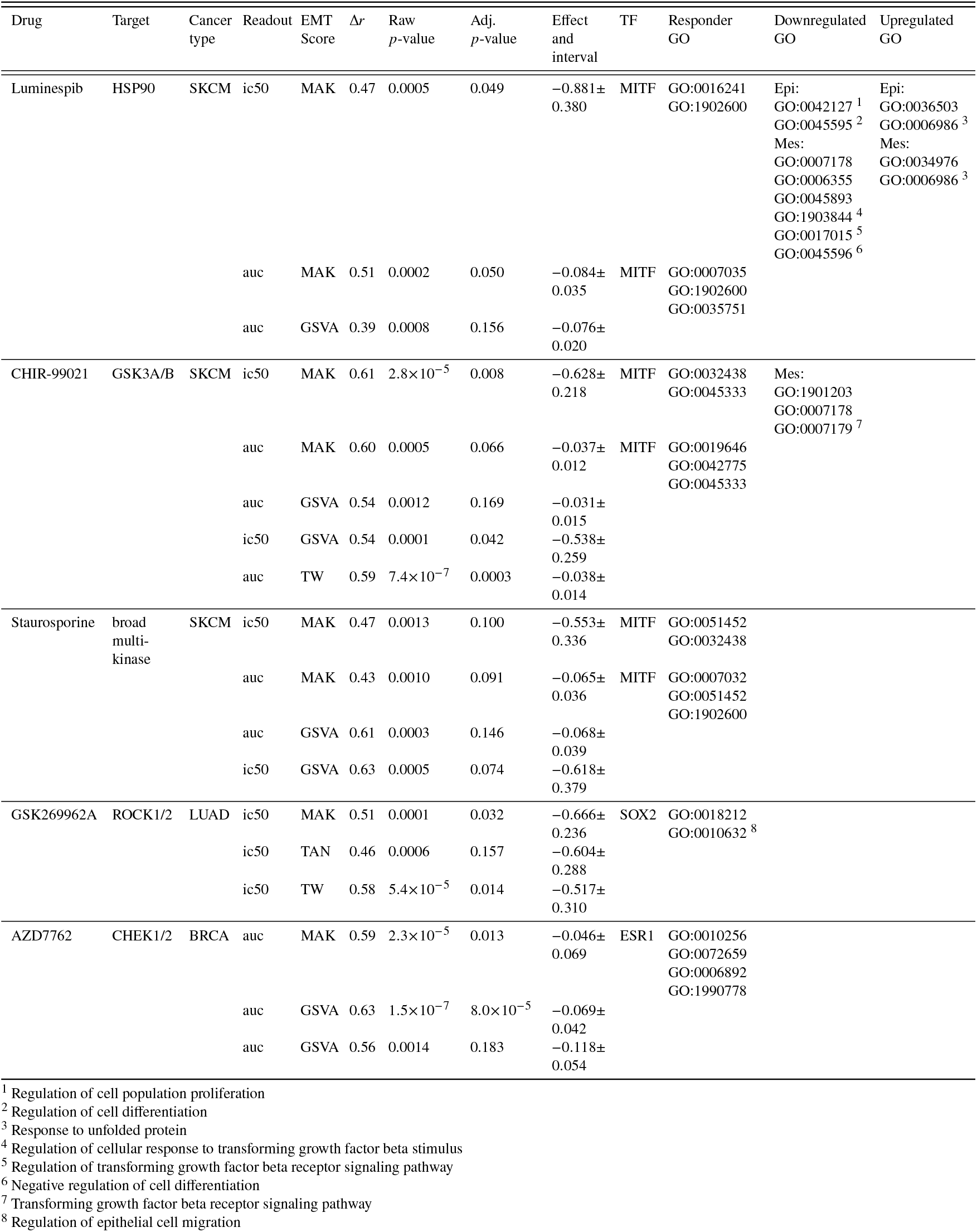
The five top-ranked EMT-dependent compounds. Each of the five compounds is characterised by its name, target and cancer type for which the association was found. The statistics for the ablation study (Δr and (adjusted) *p*-value) and causal inference (effect size plus interval in terms of Δlog(IC_50_) or ΔAUC) are given for each response readout and EMT score. Furthermore, the enriched TFs and GO terms for the responding cell lines and the enriched GO terms in transcriptional signatures for the compounds in SKCM are shown. Selected GO terms are annotated in the footnotes.

**Figure 3:**
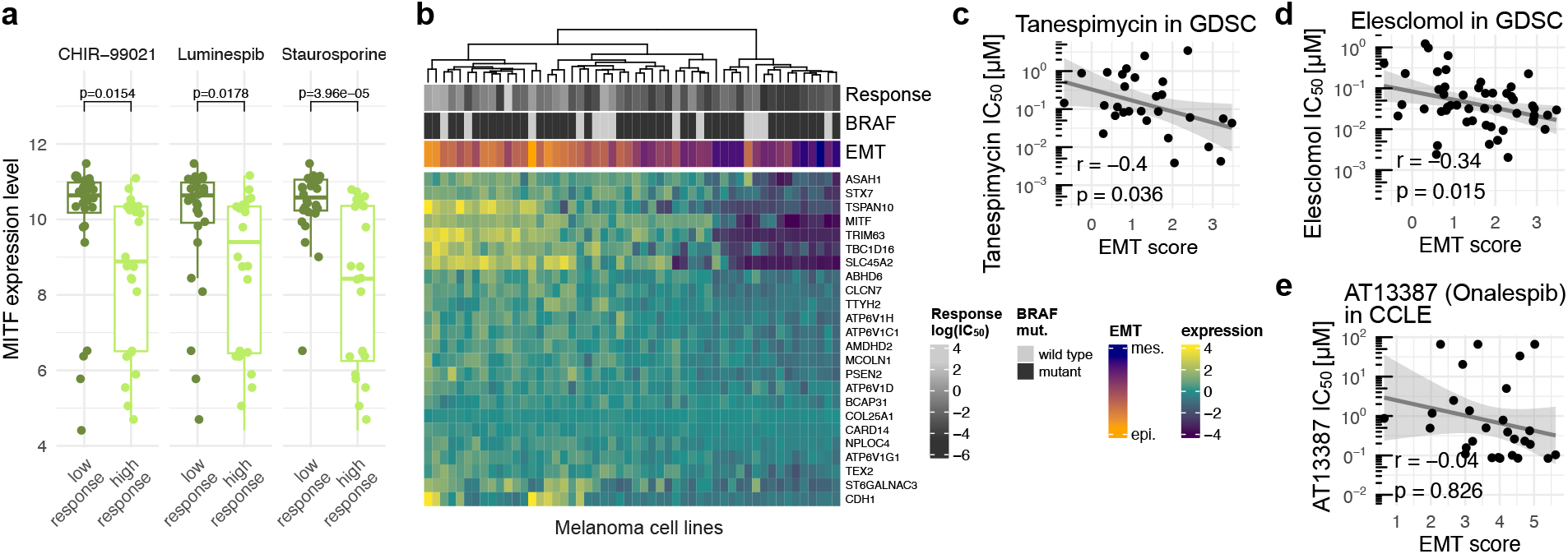
EMT is associated with transcription factors and susceptibility to HSP90 inhibitors. (**a**) Boxplots of binarised drug response (i.e. discretisation by median IC_50_ values) of CHIR-99021, luminespib and staurosporine show associations with *MITF* expression levels quantified by a two-sided *t*-test and its derived *p*-value. The centre on the boxplot represents the median, while the box illustrates the interquartile range (IQR). The whiskers show a range that is 1.5 times the size of the IQR. (**b**) A heatmap shows differentially expressed genes of luminespib drug response quantified by log(IC_50_) values in SKCM cell lines (FDR < 0.1) for the subset of MITF targets. Scatter plots show drug susceptibility of SKCM cell lines to other HSP90 inhibitors such as (**c**) tanespimycin, (**d**) elesclomol and (**e**) AT13387 (onalespib) in an independent dataset. The solid line depicts a fitted ordinary least squares regression model with its 95% confidence interval. The Pearson’s correlation coefficient (r) and the associated *p*-value of the correlation test (p) are displayed.

To interpret the five top-ranked EMT-dependent compounds, we employed gene set enrichment analysis of the set of differentially expressed genes between higher and lower responding cell lines leveraging the entire transcriptome (**Supplementary Data 3**; **Methods**). The enrichment analysis found upstream transcription factors (TF) acting as master regulators that can affect both molecular EMT markers and the set of differentially expressed genes. For example, we found that responses to the CHK1 inhibitor AZD7762 in BRCA were successfully predicted by the MAK and GSVA EMT scores (**Fig. 2e**; **Supplementary Fig. 3c**; **Table 1**), and identified that the TF target genes of ESR1 were enriched (adj. *p* = 4.3 ×10^−28^, odds ratio = 2.44, Fisher’s exact test; **Supplementary Data 3**; **Table 1**). Furthermore, lower response to AZD7762 was associated with higher *ESR1* expression (**Supplementary Fig. 4a**).

*ESR1* expression is associated with clinical BRCA subtypes (PAM50), especially the invasive basal BRCA subtype is characterised by low *ESR1* expression ^29^. Accordingly, we confirmed that cell lines derived from the more invasive basal-like BRCA displayed higher MAK EMT scores resembling the mesenchymal phenotype (*p* = 0.002, ANOVA F-test; **Fig. 2g**). Therefore, we added the PAM50 subtype to the EMT score and *ESR1* expression as fixed effects in a regression model predicting AZD7762 response and found that it did not further improve our model (*p* = 0.64, ANOVA F-test for multiple regression coefficient; **Fig. 2g**). Similar to *ESR1* expression, it is established that BRCA1/2 regulates the cell cycle by activating CHK1 in response to DNA damage and its mutations are associated with oncogenesis ^30^. Thus, we repeated the same analysis by excluding cell lines that carry *BRCA1/2* mutations (*p* = 0.73, ANOVA F-test for multiple regression coefficient; **Fig. 2g**), which also did not further improve our model. Concordantly, EMT regulators were previously shown to underlie DNA damage responses through their interaction with CHK1/2 (target of AZD7762) in BRCA cells ^31^. In summary, EMT as a predictive biomarker for AZD7762 response in BRCA reflected but was not further enhanced by BRCA subtypes and somatic mutations in *BRCA1/2*.

Furthermore, we observed performance gains for the ROCK1 (Rho kinase 1) inhibitor GSK269962A, to which lung adenocarcinoma mesenchymal-like cell lines with a higher MAK, TAN and TW EMT score were more responsive (**Supplementary Fig. 4b**; **Table 1**). We identified an associated TF SOX2 (adj. *p* = 1.5 ×10^−13^, odds ratio = 8.95, Fisher’s exact test; **Supplementary Data 3**; **Table 1**), which was previously found to be associated with EMT and metastasis in multiple cancer types, including lung cancer ^32^. We expanded the enrichment analysis of the set of differentially expressed genes for responder cell lines to Gene Ontology (GO) biological processes and found that upregulated genes in LUAD cell lines responding to GSK269962A were enriched in genes involved in the regulation of epithelial cell migration (adj. *p* = 0.0004, odds ratio = 35.07, Fisher’s exact test; **Supplementary Data 4**; **Table 1**; **Methods**), which is orchestrated by ROCK1 ^33^.

In summary, our proposed method was able to robustly identify compounds in HTS that demonstrated distinct drug responses depending on EMT, its upstream regulators and related processes in several cancer types. In the next section, we focused on elucidating further mechanisms on the compounds identified in SKCM.

### Potential regulators and drug response mechanisms of EMT in melanoma cell lines

We focused on the three remaining compounds in SKCM, i.e. CHIR-99021, luminespib and staurosporine, for which mesenchymal-like SKCM cell lines showed higher sensitivity consistently across at least three models (**Fig 2e,f**; **Table 1**). In the TF enrichment analysis (**Methods**), we found that MITF was enriched in the set of differentially expressed genes for these compounds, i.e. CHIR-99021 (adj. *p* = 3.7 ×10^−36^, odds ratio = 3.44), luminespib (adj. *p* = 6.8 ×10^−7^, odds ratio = 7.54) and staurosporine (adj. *p* = 5.3 ×10^−89^, odds ratio = 4.47; Fisher’s exact test; **Supplementary Data 3**; **Table 1**) and showed responses associated with *MITF* expression (**Fig. 3a**). For luminespib, 39 genes were significantly downregulated in responding mesenchymal-like cells, from which 23 were putative MITF target genes (**Fig. 3b**). MITF is a melanocyte master regulator and is often described as an oncogene in melanoma ^34^. It was proposed to act as a phenotype-switching regulator in melanoma, for which cells with trace MITF levels show senescent properties characterised by cell cycle arrest and cell motility, low-to-intermediate MITF levels display proliferative properties, and higher MITF levels can drive cell differentiation ^35–38^.

The MAK EMT score in SKCM was associated with previously proposed SKCM subtypes ^39^ (*p* = 5.6 ×10^−8^, ANOVA F-test; **Fig. S4c**-**e**), i.e. melanocytic cell lines characterised by high *MITF* expression showed low EMT scores (**Fig. S4c**-**e**). To quantify their impact on responses to the three compounds, we added these SKCM subtypes to the MAK EMT score and *MITF* expression as fixed effects in a regression model predicting IC_50_ values. Modelling subtypes improved predictions for staurosporine (*p* = 0.0008, ANOVA F-test for multiple regression coefficient; **Fig. S4c**), whilst we did not observe improvements for luminespib (*p* = 0.31, ANOVA F-test for multiple regression coefficient; **Fig. S4d**) or CHIR-99021 (*p* = 0.24, ANOVA F-test for multiple regression coefficient; **Fig. S4e**), thus highlighting the predictive capability of EMT in SKCM.

To gain further insights into the mechanisms of luminespib, CHIR-99021 and staurosporine, we extracted transcriptional signatures from the Library of Integrated Network-Based Cellular Signatures (LINCS) ^40^. We retrieved luminespib signatures of mesenchymal-like A375 and epithelial-like SK-MEL-28 SKCM cell lines and tested the 100 up- and down-regulated genes for enrichment in Gene Ontology (GO) biological processes (**Supplementary Data 5**; **Table 1**; **Methods**). The top process for both cells was the upregulation of genes involved in the response to unfolded proteins (A375: adj. *p* = 4.9 ×10^-19^, odds ratio = 130.34; SK-MEL-28: adj. *p* = 6.0 ×10^−11^, odds ratio = 68.09, Fisher’s exact test; **Supplementary Data 5**; **Table 1**). Notably, genes involved in the regulation of TGF-β receptor signalling, such as *SMAD3*, were significantly downregulated among the top two enriched processes (A375: adj. *p* = 0.0004, odds ratio = 17.55, Fisher’s exact test; SK-MEL-28: adj. *p* = 0.02, odds ratio = 8.81, Fisher’s exact test; **Supplementary Data 5**; **Table 1**), suggesting that luminespib response may depend on TGF-β signalling components. Commonly downregulated genes of the CHIR-99021 signature included *SMAD3* and *PXN*, which were also enriched in TGF-β receptor signalling (A375: adj. *p* = 0.0006, odds ratio = 123.73, Fisher’s exact test; **Supplementary Data 5**; **Table 1**), whereas for the staurosporine signature, TGF-β receptor signalling showed trends of enrichment (A375: adj. *p* = 0.06, odds ratio = 23.7, Fisher’s exact test; **Supplementary Data 5**).

In summary, these results demonstrated that response to the HSP90 inhibitor luminespib in the GDSC HTS may depend on EMT components, their regulators MITF and TGF-β, and their associated subtypes. Therefore, the next section assessed the generalisation of EMT-dependent drug responses to other HSP90 inhibitors and their validation in independent HTS experiments.

### EMT is associated with drug sensitivity to HSP90 inhibition in melanoma cell lines

For evaluating the robustness of EMT as a drug sensitivity biomarker to HSP90 inhibitors in SKCM, we tested for correlations between EMT scores and sensitivity to five HSP90 inhibitors across two high-throughput drug screens (**Methods; Supplementary Fig 5**). First, we assessed the IC_50_ values of HSP90 inhibitors in the GDSC, here exemplified with tanespimycin (r = -0.40, *p* = 0.036, correlation test; cell lines with higher than mean *NQO1* expression ^41^; **Fig. 3c**), elesclomol (r = -0.34, *p* = 0.015, correlation test; **Fig. 3d**), a luminespib replicate screened in both GDSC1 and GDSC2 (r = -0.33, *p* = 0.021, correlation test; **Supplementary Fig. 5a**) and SNX 2112 (r = -0.21, *p* = 0.14, correlation test; **Supplementary Fig. 5c**). Furthermore, consistent correlations were observed for AUC values across these HSP90 inhibitors in the GDSC (**Supplementary Fig. 5a**-**h**), thus highlighting the robustness of the association between EMT and responses to HSP90 inhibition regardless of the drug response readout.

To gain further evidence across independent datasets, we calculated the MAK EMT score based on gene expression data obtained from the Cancer Cell Line Encyclopaedia (CCLE) ^42^ and analysed the HTS of the Cancer Therapeutics Response Portal (CTRP) ^43^ (**Supplementary Data 6**; **Supplementary Fig 5i**-**m**). The AUC values of the screened HSP90 inhibitors SNX 2112 (r = -0.44, *p* = 0.002, correlation test; **Supplementary Fig. 5j**) and tanespimycin (r = -0.47, *p* = 0.001, correlation test; **Supplementary Fig. 5l**) were significantly associated with the EMT score in this independent HTS, and AT13387 (onalespib) displayed consistent trends (r = -0.24, *p* = 0.221, correlation test; **Supplementary Fig. 5m**).

In essence, EMT scores were consistently associated with drug sensitivity to HSP90 inhibition in SKCM cell lines across independent drug HTS and transcriptomic profiles (**Fig. 3e**; **Supplementary Fig. 5i**-**m)**. For the next section, luminespib was selected as the lead compound for further experimental validation of our method, since it showed significant performance gains with the highest estimated EMT effects (**Table 1**).

### TGF-β modulates the response to HSP90 inhibition with luminespib in epithelial-like melanoma cell lines

We conducted experiments on whether drug response to luminespib could be modulated by EMT induction. TGF-β is a known inducer of EMT ^17^, which was also suggested by the upregulation of *TGFB1* expression in many mesenchymal-like SKCM cell lines (**Supplementary Fig. 6a**-**d**). Thus, we chose two epithelial-like cell lines (IGR-37, SK-MEL-5; **Fig. 4a**) and two mesenchymal-like cell lines (RPMI-7951, A375; **Fig. 4a**), which showed different levels of sensitivity to our lead compound luminespib in the GDSC, respectively. Following a 7-day pretreatment with TGF-β1, we treated the cells with different concentrations of luminespib (**Supplementary Fig. 6e**; **Methods**) and fitted dose-response curves for each experiment to obtain IC_50_ and AUC values (**Supplementary Data 7**; **Methods**).

**Figure 4:**
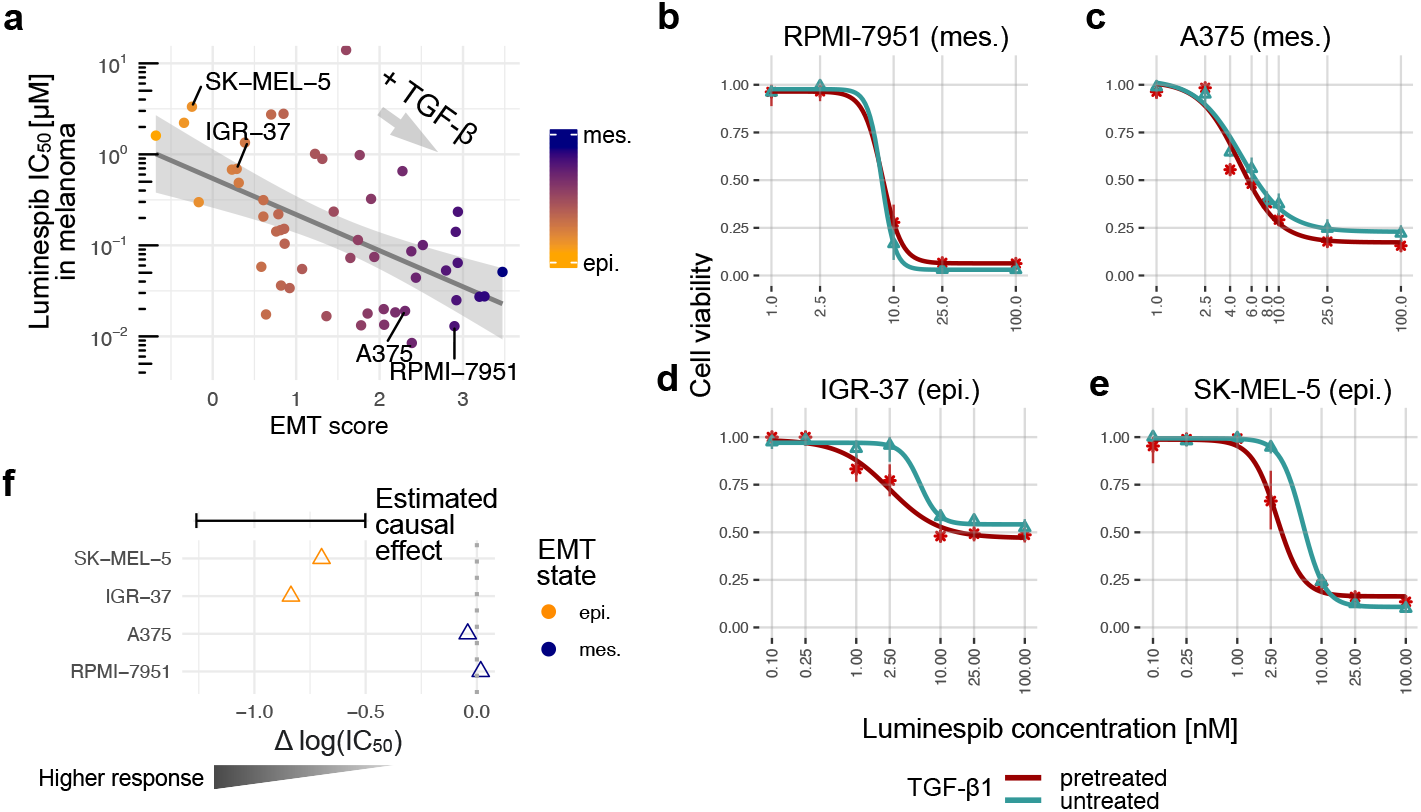
TGF-β sensitises melanoma cell lines to luminespib. (**a**) The scatter plot shows luminespib drug response stratified by EMT scores in SKCM cell lines and highlighted selected cell lines. The mesenchymal cell lines (**b**) RPMI-7951 and (**c**) A375 showed indistinguishable luminespib response upon TGF-β pretreatment. In contrast, the two epithelial cell lines (**d**) IGR-37 and (**e**) SK-MEL-5 responded stronger after pretreatment with TGF-β. Shown are the derived cell viabilities averaged across replicates and the fitted dose-response curves (**Methods**). (**f**) TGF-β treated epithelial cell lines demonstrate higher responses (decreased IC_50_) to luminespib within the predicted causal effect 95% CI.

While the mesenchymal RPMI-7951 and A375 showed no distinguishable change in luminespib response upon TGF-β1 treatment (**Fig. 4b**,**c**), the epithelial cell lines IGR-37 and SK-MEL-5 displayed increased luminespib sensitivity (**Fig. 4d,e**). To quantify this effect, we calculated the difference in log(IC_50_) values, i.e. Δlog(IC_50_), for the screened cell lines and compared it to the 95% CI of the predicted causal effect upon change in the EMT score (**Methods**). Accordingly, the epithelial cell lines IGR-37 and SK-MEL-5 showed decreased IC_50_ within this CI (**Fig. 4f**). Analogously, we compared differences in AUC values (ΔAUC), which showed consistency within the CI of the predicted causal effect (**Supplementary Fig. 6f**). In summary, this highlights that EMT can modulate HSP90 inhibitor response in epithelial-like SKCM cell lines.

## Discussion

We presented a drug response analysis encompassing the causal exploration of EMT in the context of mutational backgrounds and their upstream regulators and processes. We quantified EMT based on molecular biomarkers from gene expression profiles, thus offering a continuous score that accounts for the spectrum of intermediate and hybrid EMT states. By combining predictive and causal modelling, we identified five compounds across three cancer types with robust associations across different EMT scoring methods and drug response readouts (**Table 1**). Exemplifying our approach, we found that mesenchymal-like cell lines showed increased sensitivity to HSP90 inhibitors, particularly luminespib, which we experimentally validated.

Our pharmacogenomic modelling approaches corrected for confounders from the mutational background. Therefore, the estimated EMT effects from the causal modelling approach assumed no hidden confounders in the gene expression data. In order to address this, we performed *post hoc* differential gene expression analyses considering all genes to identify transcription factors as upstream regulators and GO biological processes. Furthermore, we mined drug transcriptional signatures to identify transcriptional confounders. Our analysis pursued the contribution of EMT on drug responses, however our systematic and causal modelling framework is generalisable to any putative drug response biomarker and its mechanisms.

We showed that epithelial-like cell lines can become more responsive to luminespib upon TGF-β treatment, whereas mesenchymal-like cell lines displayed no distinguishable change in their response. In addition, drug responses to luminespib quantified by IC_50_ values in non-TGF-β treated cells were comparable to values observed in the GDSC. Therefore, the sensitivity of some melanoma cell lines to luminespib may indeed be induced by a phenotypic conversion of cell lines. Although molecular markers do not fully capture all intricacies of the EMT program, the sensitisation of epithelial-like cell lines upon TGF-β treatment to luminespib suggests that EMT markers with their regulator MITF in melanoma may constitute a promising biomarker for selectively targeting epithelial-mesenchymal transitioning cells.

HSP90 is an ATP-dependent molecular chaperone necessary for protein folding and stabilisation of oncogenic proteins including BRAF and TGF-β receptors ^44,45^. In melanoma, HSP90 levels have been found to correlate with melanoma progression metrics such as Breslow’s depth and Clark level ^46^. The effect of HSP90 inhibition on cell viability seems to depend on MITF, which is a master regulator in melanoma cells that allow phenotype switching between distinct states ranging from arrested to highly invasive or highly proliferative phenotypes ^34,37,38^. TGF-β induces EMT across many cancer entities ^47^, and has inhibitory downstream effects on *MITF* expression ^48,49^. The sensitisation of epithelial cells via pretreatment with TGF-β suggests that TGF-β might regulate MITF in epithelial cells to allow switching to an invasive state, thereby rendering them more vulnerable to luminespib.

The exact mechanisms through which mesenchymal-like melanoma cell lines respond better to HSP90 inhibition remain elusive. They may be revealed by considering common mechanisms between the two compounds that were identified by our framework in conjunction with luminespib, namely the GSK3β inhibitor CHIR-99021 and secondly, the non-selective multi-kinase inhibitor staurosporine. Potentially, the downregulation of TGF-β signalling might be the common link between these inhibitors.

In conclusion, we demonstrated that the pharmacogenomic assessment of EMT markers with predictive and causal modelling can predict drug susceptibilities and reveal relevant tumour biology in cancer cell lines. We anticipate that considering additional parameters of EMT-like phenotype transitions, such as cell morphology and proteomics, will increase mechanistic insights to EMT and its impact on drug responses. These and other types of follow-up studies may ultimately enable the selective targeting of transitioned cancer cells from the primary tumour or circulating tumour cells to prevent dissemination and metastasis.

## Methods

### Drug response data

The drug response data from the Genomics of Drug Sensitivity in Cancer (GDSC) was obtained from its release 8.4 under https://ftp.sanger.ac.uk/project/cancerrxgene/releases/. Both GDSC1 and GDSC2 datasets were used in this analysis, using the half maximal inhibitory concentration log(IC_50_) and area under the curve (AUC) as metrics for quantifying drug responses. This resulted in 700 drug response profiles from 544 unique compounds. The Cancer Therapeutic Response Portal (CTRP) drug response data was downloaded from DepMap (https://depmap.org/portal/) contained in the file ‘CTRPv2.0_2015_ctd2_ExpandedDataset.zip’, which included 545 drug response profiles from 496 unique screened compounds.

### Somatic mutations and copy number alterations

The GDSC project has previously compiled a selection of high-confidence cancer driver genes, including somatic mutations and copy number alterations, available under http://www.cancerrxgene.org/downloads/bulk_download. These binary matrices comprised the somatic mutational status for each identified genetic event for all cancer cell lines, thus characterising their genetic landscape. They contained the status of somatic mutations from 218 cancer genes and 802 copy number segments of 775 cancer cell lines across 31 cancer types.

### Gene expression profiling and cancer subtypes

The GDSC RMA-processed Affymetrix array gene expression data was downloaded from https://www.cancerrxgene.org/gdsc1000/GDSC1000_WebResources//Data/preprocessed/Cell_line_RMA_proc_basalExp.txt.zip, containing 781 cell lines for our investigated cancer types. The CCLE log2 transformed RNA-seq gene expression data was downloaded from DepMap (https://depmap.org/portal/) contained in the file ‘OmicsExpressionProteinCodingGenesTPMLogp1.csv’ (22Q4). The BRCA (PAM50) ^29^ and SKCM ^39^ subtype annotations were obtained from the supplementary material of Jaaks *et al*. (Table S2) ^50^ and Warren *et al*. ^51^, respectively.

### EMT scores

We quantified EMT in 27 cancer types that had > 5 cancer lines available using four established methods, i.e. Mak *et al*. ^8^ (MAK), gene set variation analysis ^21^ (GSVA), Tan *et al*. ^7^ (TAN) and Tagliazucchi and Wiecek *et al*. ^10^ (TW). For the MAK EMT score, we ranked genes based on their Pearson’s correlation to four EMT marker genes, i.e. *CDH1, CDH2, VIM* and *FN1*. The genes were then ordered by their respective correlation coefficients and the top 25 genes highly correlated to *CDH1* expression were selected as ‘epithelial’ marker genes, whereas the top 25 genes that were highly correlated to each respective mesenchymal gene were grouped as ‘mesenchymal’ markers, resulting in EMT gene signatures comprised of all unique genes from the 25 epithelial and 75 mesenchymal markers for each cancer type. For each cell line, the EMT score was then calculated by the difference in mean expression levels of mesenchymal and epithelial markers.

For the GSVA EMT score, the ‘msigdbr’ R package was used to queue gene sets for the subsequent gene set variation analysis using the ‘GSVA’ R package, which yielded gene set enrichment scores from the EMT gene set for each cell line ^9^. For the TAN EMT score, we downloaded the provided tables in their supplementary material (Table S4C ^7^) and extracted the scores from their set of cancer cell lines. Similarly, for the TW EMT scores, we used the provided supplementary tables in their supplementary material to extract scores (Source data ^10^). All EMT scores are supplied in **Supplementary Data 1** and **Supplementary Data 6**.

### Predictive modelling and ablation study

Drug responses denoted by *y* and quantified by log(IC_50_) and AUC values were modelled by the EMT score *s* and somatic alterations ***x*** = {*x*_1_…., *x*_*m*_}, consisting of *m*binary encoded genetic alterations. The model was specified by

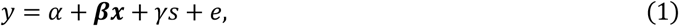

with intercept *α*, confounder coefficients ***β***, EMT coefficient *γ* and the error term *e*. The python package ‘sklearn’ was used to fit the regression model with lasso penalty and cross-validation for choosing the optimal penalty hyperparameter for each compound, cancer type and EMT score separately. For benchmarking the model performance, we performed 5-fold cross-validation with 5 repetitions. The Pearson’s correlation (r) between predicted and ground truth response was calculated on the test set to quantify model performance for each of the five folds and five initialisations.

An ablation study was conducted to prioritise drugs and cancer types for which EMT is suggested to contribute to the drug response phenotype. It was performed by refitting the models with removed EMT score *s* and recording its performance with the same splits. To compare the model performances of the full versus the model with ablation of *s*, a *t*-test for resampled performance metrics was used for assessing significant decreases of Pearson’s r across all the performances from the 25 models ^52^. The resulting p-values were corrected for multiplicity using the Benjamini-Hochberg false discovery rate (FDR) method ^53^ for each cancer type and EMT score separately. We found 32 compounds with FDR < 0.2 across six cancer types. In the main manuscript, we focused on five compounds that showed robustly significant performance differences in at least three out of eight possible models (4 EMT scores ×2 response readouts).

We only performed modelling if at least 25 cell lines for a given cancer type and drug were observed in the screening experiment. Furthermore, for modelling IC_50_ values, we did not consider models for which > 70% of IC_50_ values for a given drug and cancer type were extrapolated considerably beyond the maximum tested concentration c_max_, i.e. IC_50_ > 2c_max_. The full results are supplied in **Supplementary Data 2**.

### Causal modelling

Double machine learning (DML) is often used for estimating treatment effects on observed outcomes. It consists of two stages, (1) learning the propensity and outcome models as nuisance functions to extract their residuals, and (2) regressing outcome residuals on treatment residuals to obtain valid treatment effects and confidence intervals (CI) ^25,26^. Accordingly, we estimated the causal component of EMT by fitting a causal forest ^27^ in conjunction with DML for each drug, cancer type and EMT score, implemented in the CausalForestDML method within the python package ‘econml’ ^28^. The two nuisance functions were fitted using the same lasso regression model as used above. Thereby, we modelled the drug responses as outcome *y* to estimate the effect of the EMT score *s*as a continuous variable in the presence of the mutational background as confounders *x*. The estimated effect (EMT effect) then assesses the impact of undergoing EMT via non-mutational tumour plasticity on drug response. Since the EMT scores are continuous, the effect was given per unit of EMT change, i.e. for the interval of one standard deviation from the distribution of EMT scores for each cancer type. This effect and its 95% CI was compared with the validation experiments. The full results are supplied in **Supplementary Data 2**.

### Transcription factor and gene ontology enrichments

We sought to identify enrichments of genes correlated to drug responses in transcription factor (TF) targets and Gene Ontology (GO) biological processes from the transcriptional background of cancer cell lines. For a given drug response and transcriptomic profile within a given cancer type, we performed differential gene expression between continuous drug responses using linear models implemented in the ‘limma’ R package. The differentially expressed genes (FDR < 0.1) were then used as query genes for a gene set enrichment analysis with the ‘enrichR’ R package, for which we tested gene sets consisting of curated TF target genes ^54^ as potential upstream regulators of EMT and biological processes in the GO knowledge base ^55^. We only considered the gene set positively correlated with drug response and its top enriched TF and the top two enriched GO terms by their adjusted *p*-values including ties in **Table 1**, while the full results are supplied in **Supplementary Data 3** for TFs and **Supplementary Data 4** for GO terms.

### LINCS transcriptional signatures

The transcriptional signatures of the Library of Integrated Network-Based Cellular Signatures (LINCS) program contain sets of genes with up- and down-regulated gene expression levels upon chemical or genetic perturbations ^40^. Using the CLUE knowledge base (https://clue.io/lincs) and its provided API, we retrieved the signatures of luminespib for two SKCM cell lines, i.e. mesenchymal-like A375 and epithelial-like SK-MEL-28. We aggregated the 100 up- and down-regulated genes from all available signatures for cell lines. Then, we used these genes as a query for a gene set enrichment analysis with the ‘enrichR’ R package for each cell line to test for enrichments of GO biological processes. For staurosporine and CHIR-99021, only mesenchymal-like A375 cells were available. We used the overlapping signature genes of the transcriptional signatures of luminespib in A375 cells as a query for the same enrichment analysis in order to check for common mechanisms between the three compounds. We only considered the top two enriched GO terms by their adjusted *p*-values including ties in **Table 1**, while the full results are supplied in **Supplementary Data 5**.

### Cell culture

SK-MEL-5 (source: ATCC), A375 (source: ATCC), RPMI-7951 (source: DSMZ) were cultured in Gibco Dulbecco’s Modified Eagle Medium (DMEM) supplemented with 10% Fetal Bovine Serum (FBS) and 1% Penicillin-Streptomycin (Pen-strep) (10000 U/mL). IGR-37 (source: DSMZ) was cultured in Gibco DMEM supplemented with 15% FBS 1% Pen-strep. To induce EMT based on previous literature ^17^, the media were supplemented with 5 ng/mL TGF-β1 (R&D Systems 7754-BH/CF) for 7 days.

### Luminespib treatment

10,000 cells in 100 μL medium per well were seeded in 96-well opaque, white, flat-bottom plates. After the cells were allowed to attach at 37 ^°^C, luminespib (Selleck-Chem: S1069) dissolved in DMSO was added into the wells at the indicated concentrations and 0.5% DMSO. The negative control wells were treated with 0.5% DMSO alone, whereas the blank wells contained only the media. The plates were incubated for 72h before the CellTiter-Glo® 2.0 Cell Viability Assay (Promega: G924A) was performed following the manufacturer’s protocol. Luminescence was measured using the Perkin Elmer EnVision 2104 Multilabel Plate Reader.

### Dose-response analysis

Cell viability (*ν*) (capped between 0 and 1) was calculated with intensities from blank (*I*_*B*_), negative control (*I*_*NC*_) and luminespib treatment (*I*_*L*_) wells with

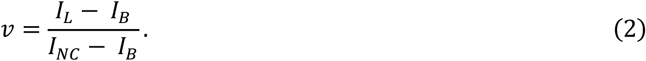

Dose-response curves were fitted and IC_50_ values were calculated using the four-parameter log-logistic (LL.4) model in the R package ‘drc’ ^56^ and AUC values were calculated using the R package ‘PharmacoGx’ ^57^. The results are supplied in **Supplementary Data 7**.

## Code accessibility

The source code for the presented analysis is available at https://github.com/mendenlab/emtpb.

## Author Contributions

Conceptualization, A.J.O. and M.P.M.; Data curation, A.J.O., G.A., M.K.S. and T.J.O.; Analysis, A.J.O., G.A. and M.K.S.; Methodology, A.J.O., G.A., M.K.S. and M.P.M.; Supervision, D.K. and M.P.M.; Visualisation, A.J.O. and G.A.; Writing original draft, A.J.O., M.K.S., G.A. and M.P.M.; Writing, review and editing, all authors.

