## Supplementary information for "The pharmacogenomic assessment of molecular epithelial-mesenchymal transition signatures reveals drug susceptibilities in cancer cell lines"

### This PDF file includes:

Supplementary Figure 1 to 6

### Other supplementary information for this manuscript include:

Supplementary Data 1 to 7

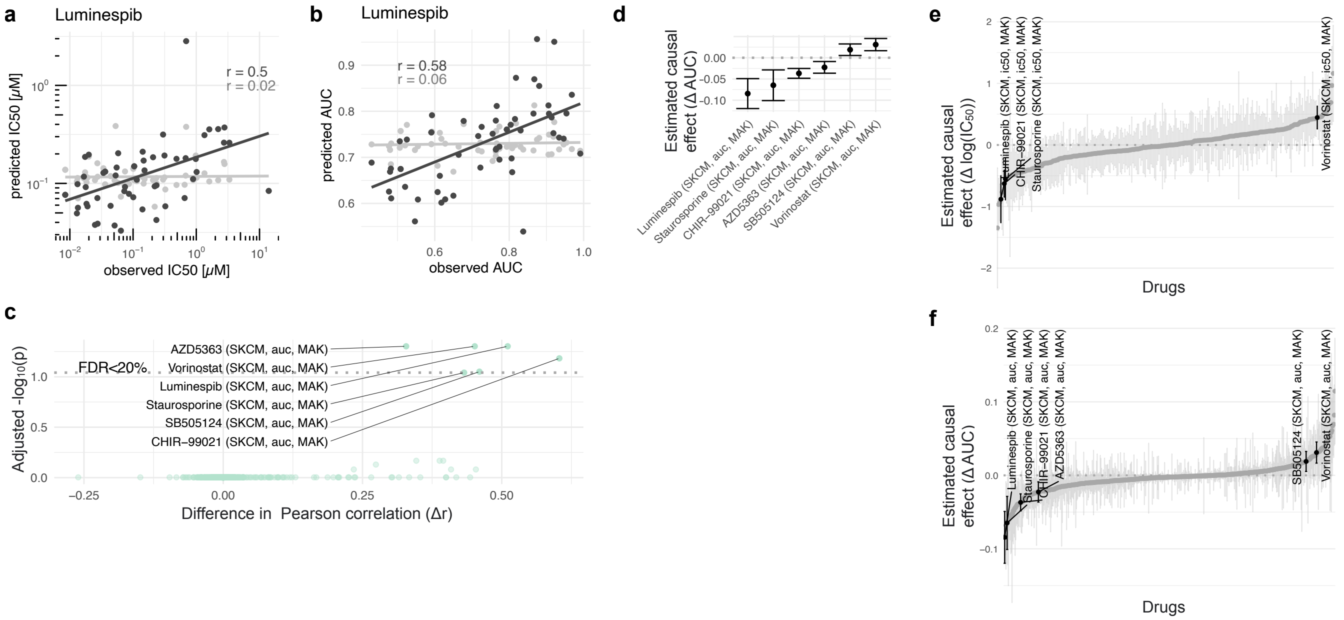


**Supplementary Figure 1: Predictive and causal modelling in SKCM.** The Scatter plot shows observed versus predicted (**a**) $IC_{50}$ and (**b**) AUC values of luminespib for the full model (black: leveraging EMT scores and mutational background) and with ablation of the MAK EMT score (grey). (**c**) The volcano plot illustrates the ablation study in SKCM with AUC quantifying significance and effect size with adjusted $p$-values of a $t$-test for performance metrics and the difference in Pearson's correlation Δr, respectively. (**d**) The inferred EMT effects and 95% confidence intervals (CI) on drug responses in terms of AUC for the identified set of compounds in SKCM are shown. Furthermore, the estimated EMT effects with 95% CIs for all compounds in SKCM using (**e**) $IC_{50}$ and (**f**) AUC values coloured in grey with significant compounds in black are highlighted.

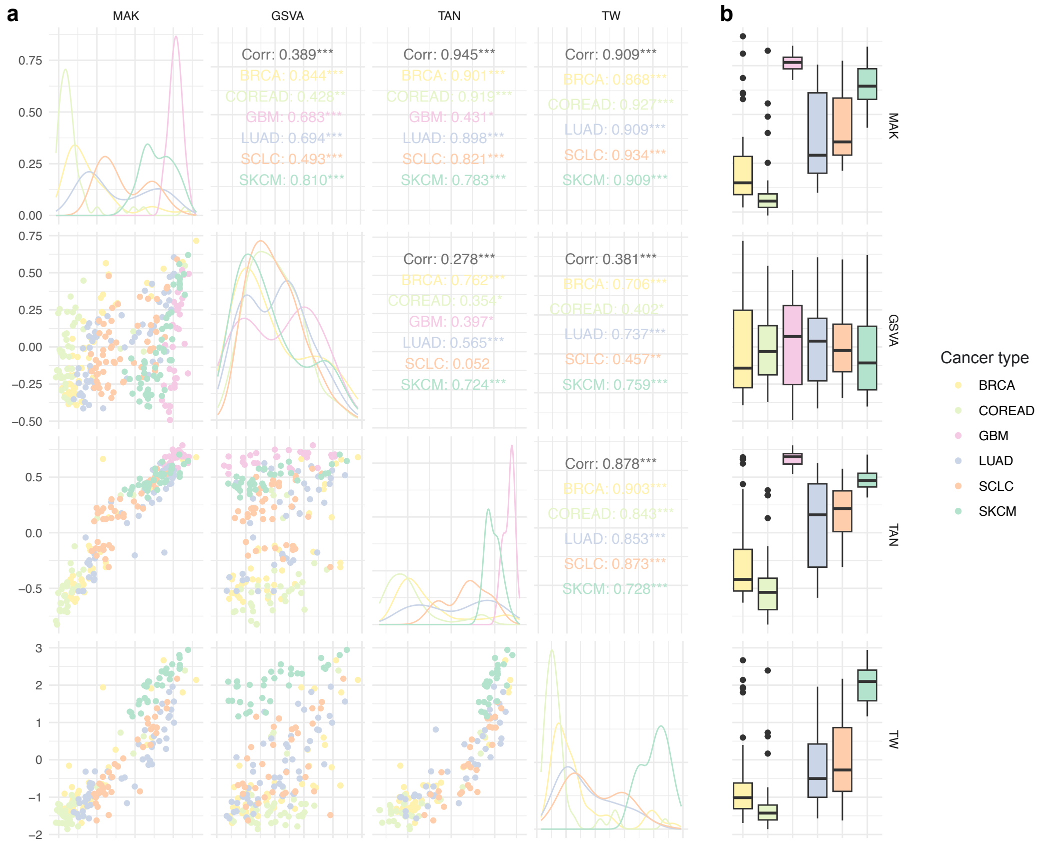


**Supplementary Figure 2: Comparison of EMT scores.** (**a**) Pairwise plots of the four derived EMT scores display their scatter plots (lower off-diagonal), their Pearson's correlation coefficient including significance of the correlation test, i.e. '***' for $p < 0.001$, '**' for $p < 0.01$, '*' for $p < 0.05$, '.' for $p < 0.1$ and no asterisk otherwise (upper off-diagonal), and their distribution (diagonal). (**b**) Boxplots demonstrating EMT scores of six cancer types that show at least one significant compound (FDR < 0.2) in the predictive modelling approach. The centre represents the median, while the box illustrates the interquartile range (IQR). The whiskers show a range that is 1.5 times the size of the IQR.

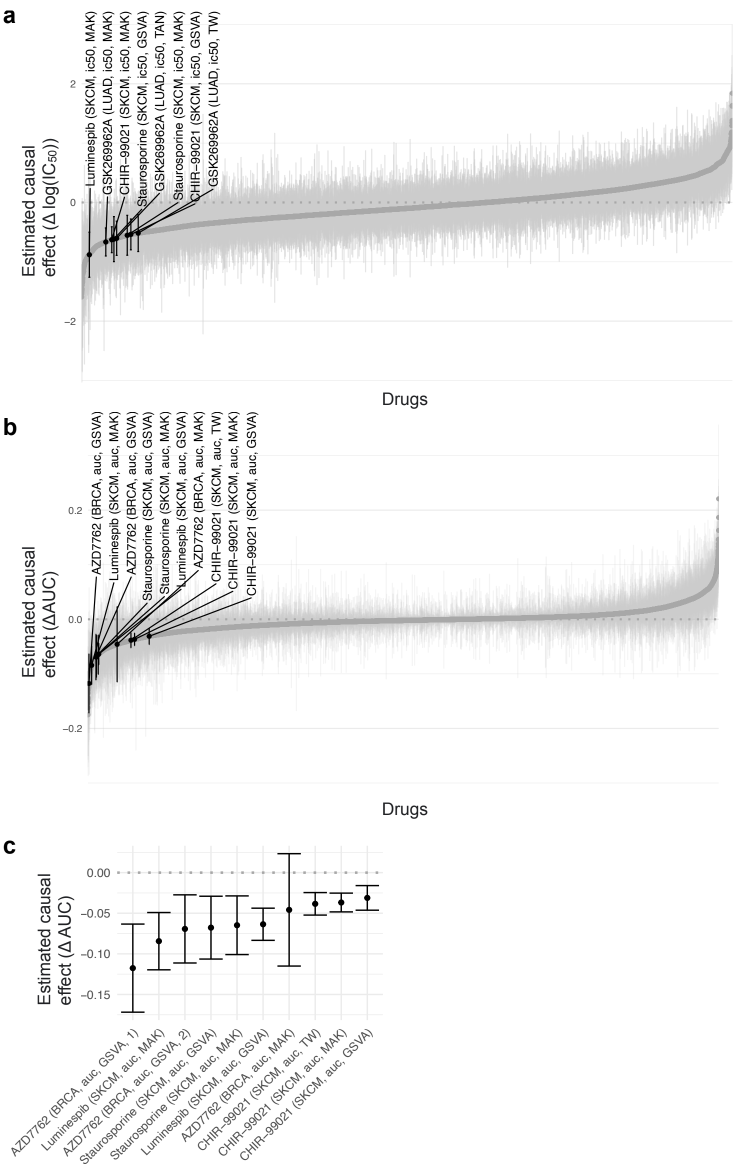


**Supplementary Figure 3: Causal modelling across cancer types.** The estimated EMT effects with 95% confidence intervals (CI) for all compounds across all cancer types using (**a**) $IC_{50}$ and (**b**) AUC values coloured in grey with significant compounds in black are highlighted. (**c**) Furthermore, models for AUC values across all cancer types and EMT scores, the estimated EMT effect plus 95% CIs for the significant compounds are shown.

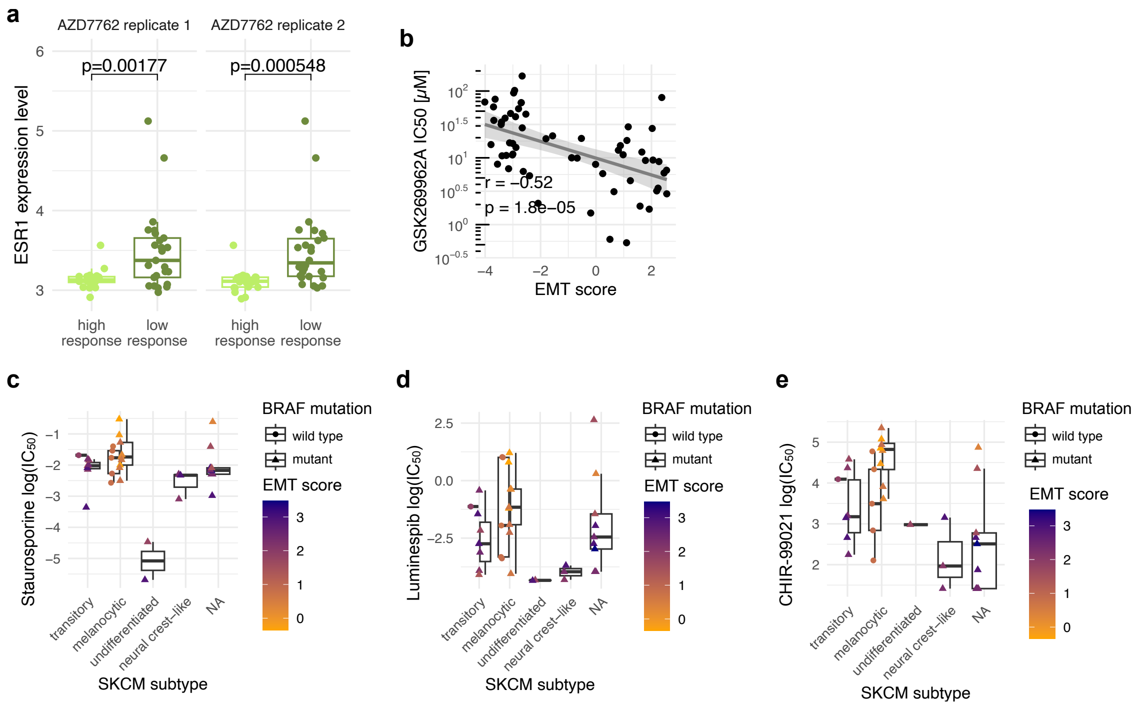


**Supplementary Figure 4: Characterisation of EMT in the context of drug response.** (**a**) The boxplots show *ESR1* expression of BRCA cell lines stratified by higher versus lower AZD7762 responding cell lines, i.e. discretisation by median $IC_{50}$ quantified by a two-sided $t$-test and its derived $p$-value. (**b**) The scatter plot highlights $IC_{50}$ values of LUAD cell lines treated with the ROCK inhibitor GSK269962A dependent on their MAK EMT score. The Pearson's correlation coefficient (r) and the associated $p$-value of the correlation test (p) are displayed. The boxplots show the response to (**c**) staurosporine, (**d**) luminespib and (**e**) CHIR-99021 depending on BRCA PAM50 subtypes, the MAK EMT score and mutations in BRCA1/2. The centre on the boxplot represents the median, while the box illustrates the interquartile range (IQR). The whiskers show a range that is 1.5 times the size of the IQR.

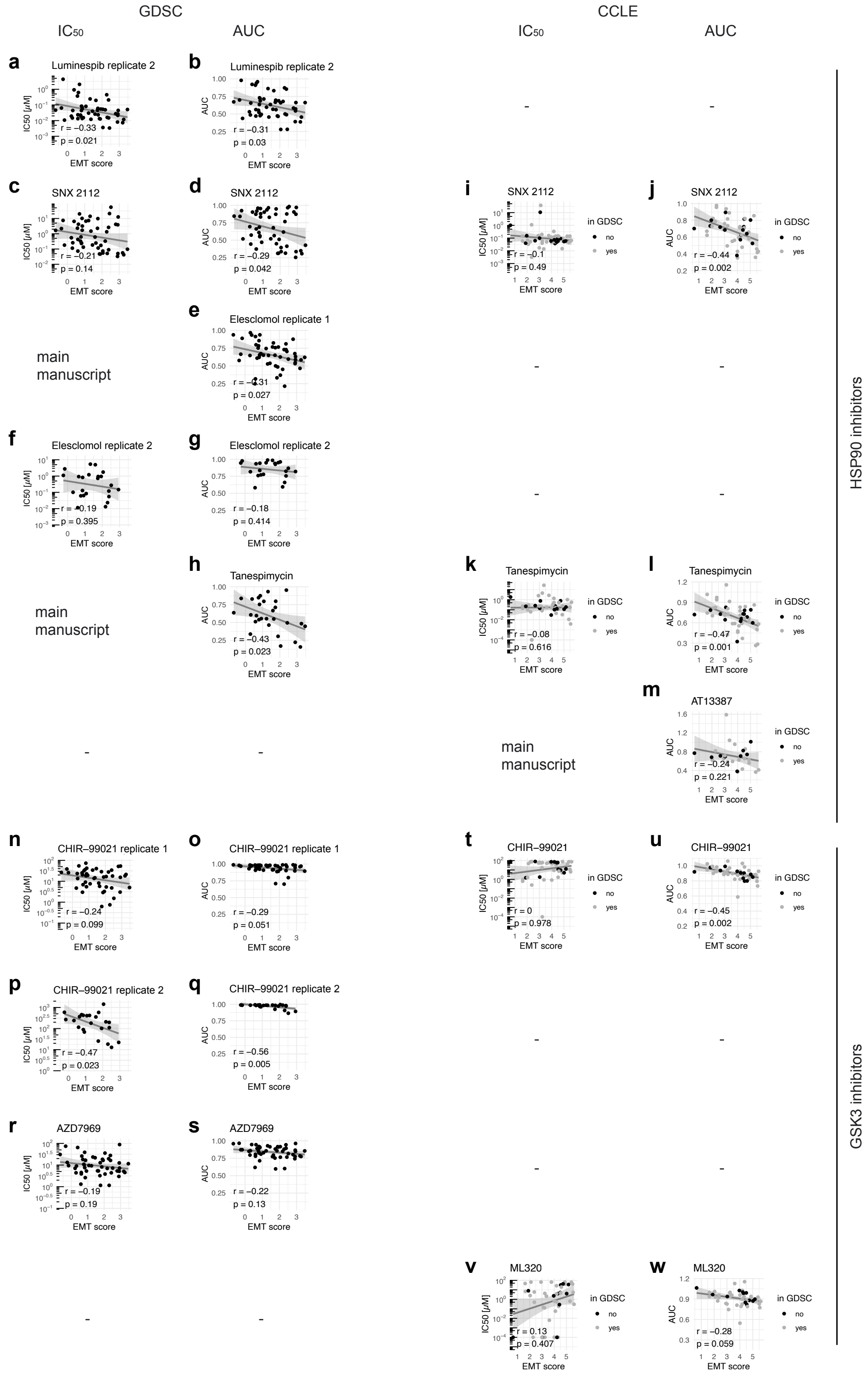


**Supplementary Figure 5: Consistency of HSP90 and GSK3 inhibitors in GDSC and CCLE.** (**a-h**) The scatter plots show drug responses to HSP90 inhibitors in the GDSC quantified by either $IC_{50}$ or AUC and their MAK EMT scores for the second luminespib replicate, SNX 2212, both elesclomol replicates and tanespimycin. The Pearson's correlation coefficient (r) and the associated *p*-value of the correlation test (p) are displayed. (**i-m**) The analogous scatter plots show HSP90 inhibitors SNX 2112, tanespimycin and AT13387 (onalespib) in the CCLE/CTRP. Cell lines in grey were also screened in the GDSC datasets, whereas cell lines in black were exclusively screened in the CCLE/CTRP dataset. (**n-s**) The scatter plots show drug responses to GSK3 inhibitors in the GDSC quantified by either $IC_{50}$ or AUC and their MAK EMT scores for the two CHIR-99021 replicates and AZD7969. (**t-w**) The analogous scatter plots show GSK3 inhibitors CHIR-99021 and ML320 in the CCLE/CTRP. Cell lines in grey were also screened in the GDSC datasets, while cell lines in black were exclusively screened in the CCLE/CTRP dataset. Dashes in the shown figure indicate missing drugs in the GDSC and CCLE/CTRP datasets, respectively.

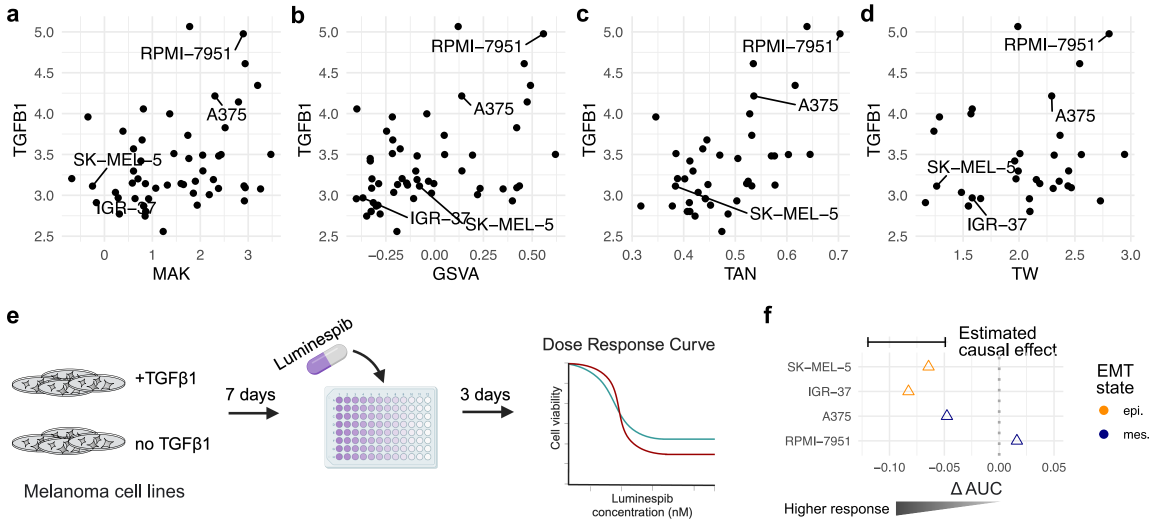


**Supplementary Figure 6: Validation experiments for luminespib with TGF-β pretreatment.** *TGFB1* expression is demonstrated in SKCM cell lines depending on their EMT scores: (**a**) MAK, (**b**) GSVA, (**c**) TAN, and (**d**) TW. (**e**) The cartoon for luminespib screens illustrates TGF-β1 pretreatment. (**f**) TGF-β treated epithelial cell lines demonstrate higher responses (decreased AUC) to luminespib within the inferred causal effect 95% confidence interval.
